# AKT signaling in hepatocytes rapidly increases glucose phosphorylation and contribution to glycogen without affecting metabolite pool sizes or glycogen breakdown

**DOI:** 10.64898/2025.12.01.691331

**Authors:** Megan Stefkovich, Won Dong Lee, Talia Coopersmith, Joseph A. Baur, Joshua D. Rabinowitz, Paul M. Titchenell

**Affiliations:** Institute of Diabetes, Obesity, and Metabolism, University of Pennsylvania, Philadelphia, Pennsylvania, USA; Department of Physiology, University of Pennsylvania, Philadelphia, Pennsylvania, USA; Lewis Sigler Institute for Integrative Genomics, Princeton University, Princeton, New Jersey, USA; Department of Chemistry, Princeton University, Princeton, New Jersey, USA; Ludwig Institute for Cancer Research, Princeton Branch, Princeton, New Jersey, USA; Department of Biochemistry, Yonsei University, Seoul, Republic of Korea; Penn College of Arts and Sciences, University of Pennsylvania, Philadelphia, Pennsylvania, USA

**Keywords:** glycogen, glucose homeostasis, glucokinase, glucose 6-phosphate, glycolysis, gluconeogenesis

## Abstract

**Background and aims:** Hepatic insulin action is essential for whole body glucose homeostasis. Insulin’s inhibition of glycogen breakdown, suppression of gluconeogenesis, and activation of glycogen synthesis are critical for postprandial glucose disposal. AKT, a serine-threonine kinase and well-established insulin signaling target, regulates hepatic glucose metabolism through transcriptional and posttranslational mechanisms. However, current knowledge about AKT’s regulation of hepatic glucose metabolism largely stems from genetic loss of function models, precluding observation of rapid, transcription-independent effects.

**Methods:** Stable isotope tracing using [U-^13^C]-glucose was coupled with pharmacological inhibition of AKT using MK-2206 in primary rat hepatocytes. Bulk metabolomics was performed on AKT knockout livers and primary rat hepatocytes treated with MK-2206. Radiolabeled glucose was used to quantify short-term changes to glycogen synthesis.

**Results:** MK-2206 treatment decreased glucose contribution to glucose 6-phosphate and uridine diphosphate glucose within minutes without significantly affecting total metabolite pool sizes or changes to glucokinase protein levels. This was accompanied by a decrease in glucose contribution to glycogen, independent of changes to glycogen breakdown or glycogen synthase phosphorylation.

**Conclusions:** Together, these results demonstrate that AKT acutely regulates glucose contribution to glycogen and its upstream precursors, suggesting a transcription-independent, glucokinase-centered mechanism for glycogen synthesis through the direct pathway.

## 1. Introduction

Insulin regulation of hepatic glucose production (HGP) is critical for postprandial glucose homeostasis. Insulin inhibits HGP via two mechanisms, one involving direct action at the liver and the other working indirectly through controlling adipose tissue lipolysis [1–3]. Insulin’s rapid suppression of HGP allows for robust liver glucose uptake [1; 2; 4-7], clearing an estimated 30-60% of blood glucose [8; 9]. This response is blunted in humans with type 2 diabetes, as demonstrated by euglycemic/hyperinsulinemic clamps [10; 11]. Moreover, metabolic disease blunts insulin’s direct action on the liver, demonstrated by hyperinsulinemic clamps performed on a high fat and high fructose-fed (HFFD) conscious dog model. Regardless of a high insulin infusion rate and adequate suppression of adipose tissue lipolysis, HFFD dogs showed decreased liver glucose uptake compared to normal chow controls [5]. While an aberrant response to insulin impacts glucose homeostasis, the mechanism by which insulin rapidly suppresses liver glucose output and stimulates uptake is not completely understood.

It is well-established that insulin directly controls hepatic glucose metabolism by signaling to the serine/threonine kinase AKT [12]. AKT is a major effector of insulin signaling required for insulin-stimulated glycogen storage [13], and it mediates insulin’s suppression of HGP in the liver [12; 13]. The prevailing mechanism as to how AKT regulates HGP is that it suppresses gene transcription of gluconeogenic genes *Pck1* and *G6pc* [14], which encode for phosphoenolpyruvate carboxykinase (PEPCK) and glucose 6-phosphatase (G6Pase), respectively. However, the slow degradation of PEPCK and G6Pase is insufficient to explain how insulin shifts hepatic glucose balance from output to uptake within minutes [4; 7]. Furthermore, *Pck1* and *G6pc* expression correlates poorly with blood glucose control and decreases well after insulin achieves suppression of HGP during a clamp [12; 15; 16]. AKT may also suppress HGP by preventing glycogen breakdown. Putatively, AKT achieves this by inhibiting a negative regulator of glycogen synthesis, glycogen synthase kinase 3 beta (GSK-3β), through phosphorylation [17]. However, mutation of the AKT-regulated phosphosites on GSK-3β do not impact insulin-stimulated glycogen synthesis in the liver [18], thus additional mechanisms must be at play in hepatocytes.

Hyperinsulinemic clamp studies have elucidated insulin’s relative control of glycogen breakdown versus gluconeogenesis during its rapid suppression of HGP. Experiments in dogs suggest that insulin predominately controls HGP by suppressing glycogenolysis [19; 20]. Two similar studies suggested an increase in glycolysis and decrease in gluconeogenesis played a role [7; 15], but gluconeogenic control depended on insulin’s anti-lipolytic effects [7]. Other studies rebut this, finding that a low dose of insulin, even when delivered via the portal vein, does not affect gluconeogenesis while achieving suppression of glycogenolysis. This was established using a 3-^3^H-glucose tracer method or by measuring ^14^C-phosphoenolpyruvate contribution to glucose 6-phosphate (G6P) and plasma glucose [19; 20]. In terms of human data, one study showed that glycogenolysis is more sensitive to insulin but that gluconeogenesis decreases as well [21]. Nevertheless, the majority of literature suggests that insulin’s inhibition of glycogenolysis is the dominant direct regulator of HGP compared to gluconeogenesis.

Current knowledge regarding insulin’s suppression of glycogenolysis invokes the concept of substrate push. Specifically, it suggests a “pull” mechanism, whereby suppression of glycogenolysis reduces the concentration of glycogenic precursors like glucose 1-phosphate and G6P. This could favor glucose contribution to these glycogen precursors, stimulating more glucose uptake and glycogenesis. Notwithstanding strong evidence for the former, it does not preclude a “push” mechanism, whereby insulin promotes glucose phosphorylation to provide eventual substrates for glycogen synthesis. Higher concentrations of glycogen precursors would then decrease the “pull” for glycogen breakdown. Beyond providing the building blocks of glycogen, higher levels of glycogenic substrates could also serve as allosteric regulators. Specifically, G6P has long been known to allosterically activate glycogen synthase [22]. Indeed, mutation of glycogen synthase’s G6P binding site reduced insulin-stimulated glycogen deposition in hepatocytes [23], suggesting that if a “push” mechanism exists, it may also involve allosteric regulation.

A potential effector of a “push” or allosteric mechanism could be glucokinase (GCK). Metabolic control analysis from hyperinsulinemic clamp data suggests that GCK is the rate-limiting step for insulin-stimulated glycogen synthesis [24]. This suggests that insulin may acutely promote glucose phosphorylation to facilitate blood glucose clearance and increase glycogen storage. Primary hepatocyte and rat clamp studies indicate that glycogen synthesis may not be the only beneficiary of insulin-stimulated glucose phosphorylation. Insulin increased total lactate [25–27], fructose bisphosphate, glyceraldehyde 3-phosphate, and pyruvate within 30 minutes [28] [29] [27], which correlated with a reduction in HGP in one case [29]. These studies suggest a relationship between insulin-stimulated glycolysis and suppression of HGP, but it is unclear whether increased flux through glycolysis suppresses HGP or vice versa.

Here we used both genetic loss-of-function models and pharmacological inhibition of hepatic AKT to understand the contribution of hepatic insulin signaling via AKT to long and short-term changes to glucose metabolism. Notably, we found that AKT promotes glucose contribution to G6P, UDP-glucose, and glycogen on a rapid time scale of minutes in primary rat hepatocytes. This occurred independently of changes to glycogenolysis, total metabolite levels, and phosphorylation of glycogen synthase at an insulin-regulated site, suggesting that insulin regulates glucose phosphorylation and utilization at a step upstream of its canonical regulation of glycogen storage. Secondary to the AKT-driven increase in glucose phosphorylation, gluconeogenic contribution to G6P was suppressed. Together, our data suggest that acute AKT signaling promotes glucose phosphorylation to drive glycogen synthesis, potentially through GCK activation, indirectly limiting gluconeogenic contribution.

## 2. Methods

### 2.1. Primary rat hepatocyte isolation

Primary rat hepatocytes were isolated from male Sprague Dawley rats between 150 and 175 g using the protocol described in Shen et al. 2012 [30] with a few modifications. Bovine serum albumin was added to William’s E complete medium (Gibco A1217601) to 1% w/v. To digest the liver, 20,000 units collagenase and 30 units elastase (Worthington CLSH LK002067) in addition to 1,000 Kunitz units of DNase (Worthington D2 LK003170) were dissolved in 250 mL Krebs-Ringer solution (Millipore Sigma #K4002) supplemented with 500 µM CaCl_2_ and 20 mM HEPES. All buffers and perfusion tubing were warmed to between 37 and 55°C to ensure adequate digestion. After digestion of the liver with all 250 mL of collagenase/elastase buffer, liver was dissected free and gently scraped in 150 mL of cold complete William’s E media. Cells were plated at a density of 175,000-500,000 cells per mL in 6-well tissue-culture treated plates coated with 1 mg/mL collagen in 0.2 N acetic acid. Plates were rinsed with PBS after collagen treatment and allowed to dry before plating cells. Hepatocytes were incubated for four hours in complete William’s E medium and media was changed to serum and insulin-free DMEM lacking glucose, L-glutamine, Phenol Red or sodium pyruvate (Gibco A1443001). Hepatocytes were incubated overnight for 16 hours before experiments. Experiments were never performed more than 24 hours after hepatocytes were plated.

### 2.2. [U-^13^C]-glucose isotope tracing

After overnight incubation in serum and glucose-free DMEM (Gibco A1443001), primary hepatocytes were pretreated for 30 minutes with 10 µM MK-2206 Dihydrochloride (Toronto Research Chemicals M425025), 5 µM glycogen phosphorylase inhibitor (Cayman Chemical Company 17578) or dimethyl sulfoxide (DMSO, Sigma D2650) where indicated without changing the overnight media. Addition of inhibitors to wells was staggered such that each treatment group was pre-treated for exactly 30 minutes. Cells were then treated with 10 mM [U-^13^C]-glucose and inhibitor or DMSO for the indicated amount of time. The [U-^13^C]-glucose was dissolved in the same media the cells were incubated in overnight. Media was collected 45 seconds, 1.25 minutes, 5 minutes, 10 minutes, 30 minutes or 60 minutes after the addition of the isotopically labeled glucose. Wells were washed once with ice-cold PBS and cell extraction buffer was added to the wells 1.5, 2, 5.75, 10.75, 30.75 or 60.75 minutes after addition of the isotopically labeled glucose.

### 2.3. Cell metabolite extraction

Media (200 µL) was removed from each well and added to a plastic tube containing ice-cold media extraction buffer (50:50 v/v acetonitrile:methanol) and snap frozen in liquid nitrogen. Remaining media was aspirated, and cells were washed once with ice-cold PBS. 250 µl of ice-cold cell extraction buffer (40:40:20 v/v/v methanol:acetonitrile:water) was added to each well, cells were scraped and added to a pre-chilled plastic tube and snap frozen in liquid nitrogen. Lysate was then thawed on ice and spun down at 17,000 g for 10 minutes, and supernatant was collected for LC-MS analysis.

### 2.4. Tissue metabolite extraction

Following cervical dislocation, blood was collected by cutting the inferior vena cava and livers were dissected and freeze clamped in liquid nitrogen as fast as possible. Samples were ground using a pre-cooled mortar and pestle on dry ice. To 15 mg of tissue, 600 μL cold 80:20 (v/v) methanol:water solution was added, vortexed for 10 seconds, and centrifuged at 21,000 g for 20 minutes at 4°C. The supernatants were then transferred to plastic vials for LC-MS analysis. A procedure blank was generated identically without tissue and was used later to remove the background ions.

### 2.5. Plasma metabolite extraction

Plasma (2.5 μL) was added to 60 μL cold 25:25:10 (v/v/v) acetonitrile:methanol:water, vortexed for 10 seconds, and put on ice for at least 5 minutes. The extract was then centrifuged at 21,000 g for 20 minutes at 4°C and supernatant was transferred to tubes for LC-MS analysis. A procedure blank was generated identically without plasma, which was used later to remove the background ions.

### 2.6. Metabolite measurement by LC-MS

Metabolites identified through bulk metabolomics were analyzed as described in Titchenell et al. 2016 [12]. Metabolites identified through stable isotope tracing were analyzed using a Vanquish Horizon UHPLC System (Thermo Fisher Scientific) coupled to an Orbitrap Exploris 480 mass spectrometer (Thermo Fisher Scientific). Waters XBridge BEH Amide XP Column [particle size, 2.5 μm; 150 mm (length) × 2.1 mm (i.d.)] was used for hydrophilic interaction chromatography (HILIC) separation. Column temperature was kept at 25°C. Mobile phases A = 20 mM ammonium acetate and 22.5 mM ammonium hydroxide in 95:5 (v/v) water:acetonitrile (pH 9.45) and B = 100% acetonitrile were used for both ESI positive and negative modes. The linear gradient eluted from 90% B (0.0–2.0 minutes), 90% B to 75% B (2.0–3.0 minutes), 75% B (3.0–7.0 minutes), 75% B to 70% B (7.0 –8.0 minutes), 70% B (8.0–9.0 minutes), 70% B to 50% B (9.0–10.0 minutes), 50% B (10.0–12.0 minutes), 50% B to 25% B (12.0–13.0 minutes), 25% B (13.0 –14.0 minutes), 25% B to 0.5% B (14.0–16.0 minutes), 0.5% B (16.0–20.5 minutes), then stayed at 90% B for 4.5 minutes. The flow rate was 0.15 mL/min. The sample injection volume was 5 μL. ESI source parameters were set as follows: spray voltage, 3,200 V or −2,800 V, in positive or negative modes, respectively; sheath gas, 35 arb; aux gas, 10 arb; sweep gas, 0.5 arb; ion transfer tube temperature, 300 °C; vaporizer temperature, 35°C. LC–MS data acquisition was operated under a full-scan polarity switching mode for all samples. The full scan was set as orbitrap resolution, 120,000 at m/z 200; AGC target, 1× 107; maximum injection time, 200 ms; scan range, 60–1,000 m/z.

### 2.7. Mass spectrometry data analysis

LC-MS raw data files (.raw) were converted to mzXML format using ProteoWizard (version 3.0.20315). El-MAVEN (version 0.12.0) was used to generate a peak table containing m/z, retention time, and intensity for the peaks. Parameters for peak picking were the defaults except for the following: mass domain resolution, 5 ppm; time domain resolution, 10 scans; minimum intensity, 10,000; and minimum peak width, 5 scans. The resulting peak table was exported as a .csv file. Peak annotation of untargeted metabolomics data was performed using NetID with default parameters. For tracer experiments, isotope labeling was corrected for ^13^C natural abundances using AccuCor package. For bulk metabolomics, hierarchical clustering was performed using Pearson correlation coefficient as a similarity measure. Significantly up- or downregulated metabolites were identified using a Welch’s t-test and adjusted for multiple comparisons using the Benjamini-Hochberg procedure.

### 2.8. Western blotting

Protein lysates were prepared in a modified RIPA buffer (150 mM NaCl, 50 mM Tris, pH 7.6, 1% Triton X-100, 0.5% sodium deoxycholate and 0.1% SDS) with Phosphatase Inhibitor Cocktails 2 and 3 (Sigma-Aldrich) and Roche cOmplete Protease Inhibitor Cocktail. Briefly, media was aspirated, cells were washed thrice with ice-cold PBS, and 150 µL RIPA was added per well of a 6-well plate. Cells were scraped and collected in a plastic tube, incubated on ice for 10 minutes, and spun down at 4°C for 10 minutes at 17,000 g. Supernatant was used to quantify protein concentration using the Pierce BCA Protein Assay Kit (Thermo Scientific). Sample buffer was prepared in Laemmli buffer supplemented with beta mercaptoethanol to a final concentration of 5% and sample buffer was boiled for 5 minutes at 95°C, with the exception of the samples used to blot for G6PC, which were not heated at 95°C. 30 µg of protein was loaded per well for each western blot. The following antibodies were used for immunoblotting: p-AKT Ser473 (CST no. 4060), pan-AKT2 (CST no. 4691), HSP90 (CST no. 4874), PRAS40 (CST no. 2610), p-PRAS40 (CST no. 2997), GCK (Gift from Magnuson Lab), PCK1 (Abcam 70358), PFK2 (Bethyl Laboratories A304-286A-M), FBP1 (CST No.59172), p-GS (CST no. 3891), and GS (CST No. 3886).

### 2.9. mRNA isolation and real-time PCR

To isolate mRNA from cells, media was aspirated, plates were washed twice with ice-cold PBS, and 500 µL of TRIzol Reagent (Invitrogen 15596026) was added per well of a 6 well plate. Cells were scraped and then chloroform was added at a 1:5 ratio to the TRIzol homogenate. Cells were centrifuged and the upper layer was collected.

Isopropanol was added to the supernatant to precipitate the RNA, and then RNA was extracted from cell lysates using the RNeasy Plus kit (Qiagen). Complementary DNA was synthesized using Moloney murine leukemia virus (MulV) reverse transcriptase, and the relative expression of the genes of interest was quantified by real-time PCR using the SYBR Green dye-based assay.

### 2.10. Total glycogen content

Primary rat hepatocytes were plated at 800,000 cells/well of a 6-well plate as described in Section 2.1 and were incubated overnight in fresh complete William’s E media [30] supplemented with additional fetal bovine serum (FBS) and glucose for a total concentration of 10% FBS and 25mM glucose. After 16 hours of overnight incubation, cells were pretreated with 25 µM forskolin (FSK), 5 µM Glycogen Phosphorylase Inhibitor (GPI) or DMSO for 15 minutes. Media was removed, cells were washed once with PBS, and cells were treated with 20 mM glucose +/- 25 µM FSK or 5 µM GPI in serum and insulin-free DMEM lacking glucose, L-glutamine, Phenol Red or sodium pyruvate. Cells were allowed to incubate for four hours before media was aspirated, cells were washed twice with ice-cold PBS and then 200 µL of RIPA buffer with protease and phosphatase inhibitors was added per row of the 6-well plate. Cells were scraped and protein concentration was determined using a BCA assay. Lysates were then deproteinized by adding perchloric acid to a final concentration of 1 M to the total cell lysate volume. Samples were vortexed briefly, then incubated for five minutes before centrifuging at 13,000 rpm for two minutes at 4°C. Supernatant was transferred to a fresh tube and sample was neutralized by adding ice cold 2 M KOH to a volume equal to 34% of the total supernatant. Samples were briefly vortexed and vented to release any carbon dioxide produced. Samples were centrifuged at 13,000 rpm for 15 minutes at 4°C and supernatant was collected. To measure glycogen, 250 µL of 1 mg/mL amyloglucosidase from Aspergillus niger (Millipore Sigma #A7420) was added to 50 µL of supernatant. Samples were incubated, shaking for 3 hours at 40°C. Resulting free glycosyl units and undigested samples were assayed spectrophotometrically using a hexokinase-based glucose assay kit (Millipore Sigma #G3293). Digested sample counts were multiplied by six and undigested sample counts were subtracted to determine the glycogen concentration.

### 2.11. Glycogen synthesis

Primary rat hepatocytes were plated at 800,000 cells/well of a 6-well plate. Cells were incubated overnight in serum and glucose-free DMEM. The following day, cells were pretreated with 10 µM MK2206 with or without 5 µM GPI for 30 minutes. 1 µCi of [U-^14^C]-glucose per mL media was added to each well in fresh serum and glucose-free DMEM supplemented with 10 mM glucose. Hepatocytes were incubated for 30 minutes, then media was removed and cells were washed twice with cold PBS. Each well was scraped in 70 µL of 1 M KOH and then pooled with two more wells. Samples were collected in a plastic tube and heated to 65°C for 30 minutes. Samples were then diluted in 100 µL of saturated Na_2_SO_4_ with 900 µL of 95% ethanol and left overnight at - 20°C. The following day, samples were centrifuged at 4°C for 15 minutes at 15,000 g, and then supernatant was removed and saved for quantifying total protein. The pellet was washed with 70% ethanol followed by 95% and then 100% ethanol and then resuspended in .5 mL of water. Samples were centrifuged at 4°C for 15 minutes at 15,000 g again, supernatant was discarded and pellets were resuspended in .5 mL of 95% ethanol. 200 µL of the suspension was added to 3 mL of scintillation fluid and vortexed before scintillation counting. Counts were converted to pmol and normalized to the ^14^C incubation period and total protein.

## 3. Results

### 3.1. Chronic loss of AKT signaling impacts glucose 6-phosphate levels

Previous work by Titchenell *et al*. [12] showed that inducible AKT1 and AKT2 knockout in the liver (L-AKTDKO) reduces G6P levels, suggesting that chronic loss of AKT signaling affects glucose phosphorylation. We conducted analysis on bulk metabolomics data generated from L-AKTDKO livers [12] to explore the impact of the loss of AKT on other hepatic metabolites. Notably, despite a selective decrease of G6P, levels of other glycolytic or glycogenic metabolites were surprisingly normal (Fig.1a-b). The top pathways affected were instead related to amino acid metabolism (Fig. 1c). Liver-specific knockout of AKT reduces GCK expression, the hexokinase catalyzing glucose phosphorylation in hepatocytes. Over-expression of GCK in these livers restores G6P levels [12], suggesting that the reduced G6P is dependent on changes to glucokinase and not increased glucose 6-phosphatase (G6PC) activity. These results, obtained from a two-week knockout model, highlighted the need for better understanding of glucose metabolism in the acute absence of AKT signaling, independent of transcriptional changes.

**Figure 1.**
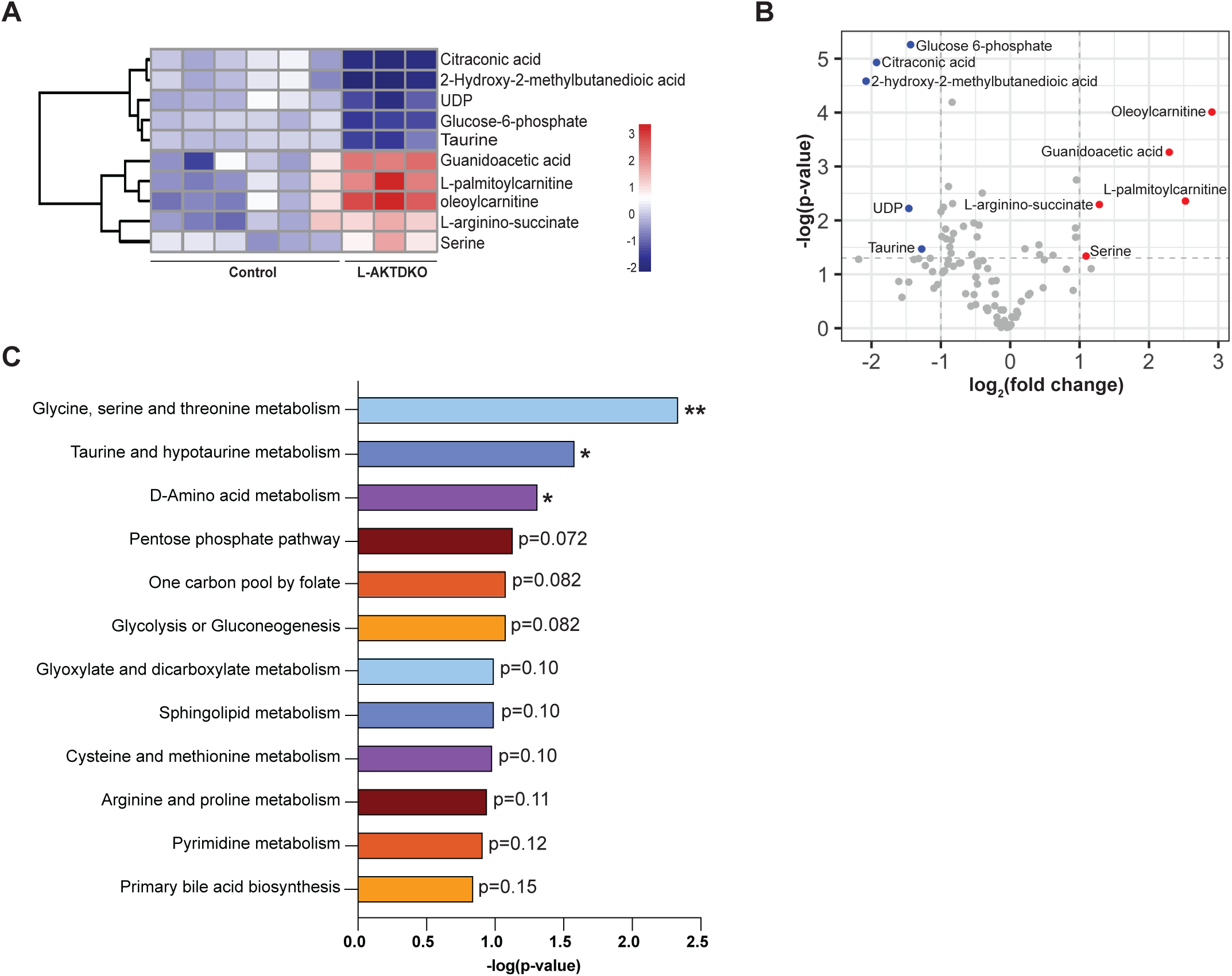
Chronic loss of AKT signaling impacts glucose 6-phosphate levels. A) Individual metabolites identified through bulk metabolomics showing metabolites significantly up- or downregulated in AKTDKO livers by an average fold-change +/- 2 with a p-value<0.05. Animals were fasted overnight for 16 hours and refed for four. Legend is log2(fold-change). B) Volcano plot and C) KEGG pathway analysis performed on bulk metabolomics described in (A). **p-value<0.005, *p-value<0.05. Enrichment determined using Fisher’s exact test. N=six control and three AKTDKO mice.

### 3.2. AKT directs glucose contribution to glucose 6-phosphate and UDP-glucose

While bulk metabolomics allows a global picture of metabolic changes, it does not provide clarity as how these changes occur. In the liver, altered glucose metabolism can be masked because a decrease in glycolytic contribution to a metabolite’s pool size can be compensated for by increased gluconeogenic or glycogenolytic contribution. For example, glycogen precursors are synthesized both directly from glucose and indirectly from gluconeogenesis (Fig. 2a). Therefore, large differences in pathway fluxes can underlie seemingly unchanged metabolite pools. Using a stable isotope tracing approach, we evaluated AKT’s acute effect on glucose metabolism with ubiquitously labeled ^13^C-glucose ([U-^13^C]-glucose) in primary rat hepatocytes (Fig 2a). An *in vitro* approach allowed us to collect samples quickly (in minutes) to determine the cell autonomous effects of AKT signaling on glucose metabolism.

**Figure 2.**
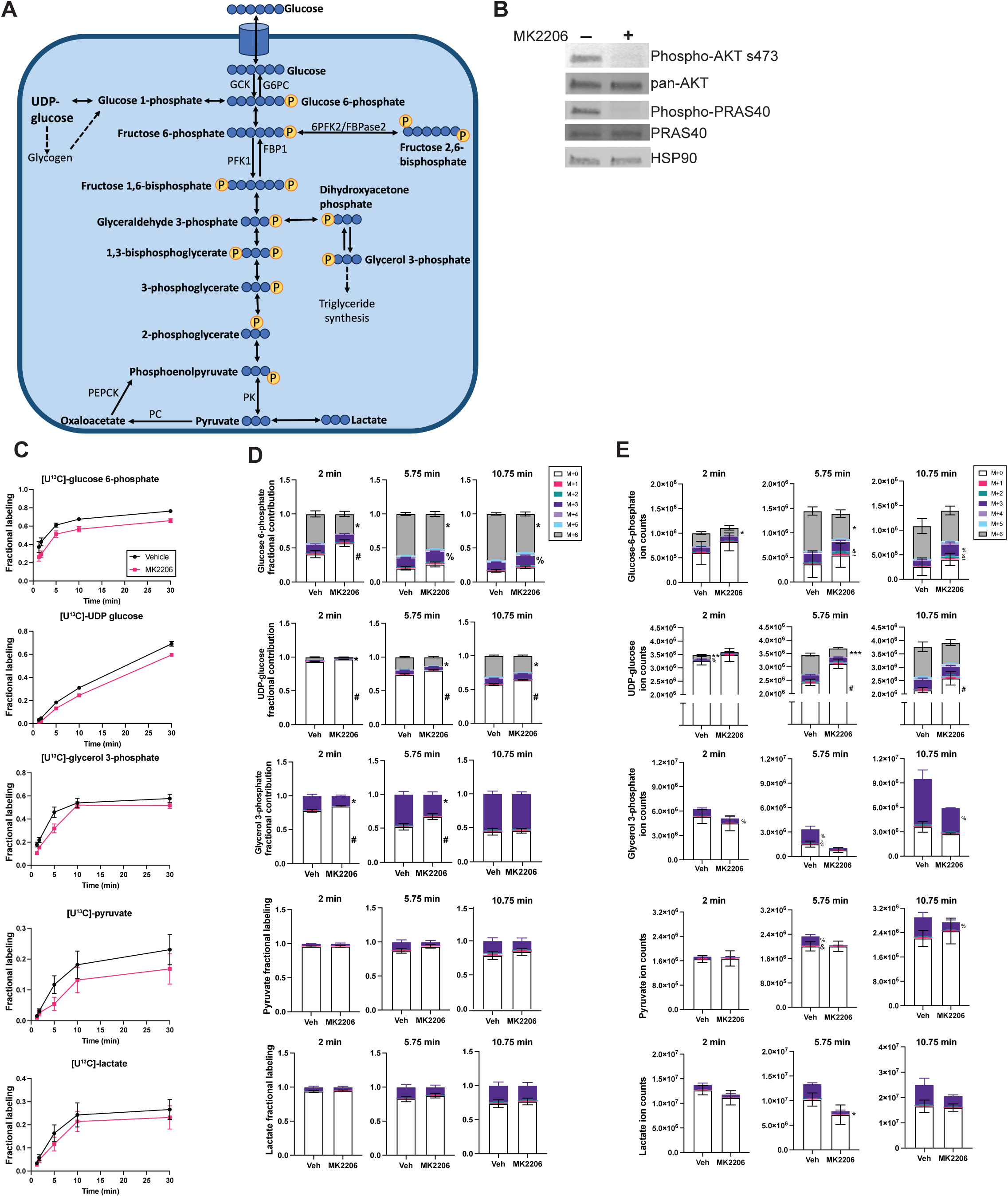
AKT directs glucose contribution to glucose 6-phosphate and UDP-glucose. A) Schematic illustrating how ubiquitously labeled ^13^C glucose contributes to glycolytic, glycogenic and lipogenic intermediates in hepatocytes.^13^C indicated by blue circles. Reversible reactions indicated by double headed arrows. Dashed arrows indicate multiple reaction steps. B) Western blot showing that the AKT inhibitor, MK2206, suppresses AKT signaling in primary rat hepatocytes. Cells were fasted overnight in serum and glucose-free DMEM, pretreated for 30 minutes with 10 μM MK2206 or DMSO in overnight media, and then treated for 30 minutes with 10 mM glucose and inhibitor or DMSO. C) Fractional labeling and D) fractional contribution of metabolite pools in primary rat hepatocytes treated with 10 mM [U-^13^C]-glucose and 10 μM MK2206 or DMSO. Cells were treated as described in Section 2.2. N=nine replicates from three pooled experiments. E) Ion counts from a representative tracing experiment included in (C) and (D). P-values calculated using Student’s t-test, values <0.05 denoted by *(M+6), % (M+3), & (M+2), ∼ (M+1), or # (M+0). P-values <0.005 or 0.0005 for M=6 denoted by ** and ***. Error bars represent +/- SEM. Time points indicate the time cell extraction buffer was added to wells.

We treated primary rat hepatocytes with a potent and selective AKT inhibitor, MK-2206 Dihydrochloride (MK2206), which effectively blocks AKT phosphorylation and that of downstream substrates like PRAS40 (Fig. 2b). Cells were incubated in glucose-free media overnight prior to a 30-minute pre-treatment with 10 µM MK2206 or DMSO vehicle control, after which experimental media was added containing 10 mM of [U-^13^C]-glucose with MK2206 or DMSO. First, we evaluated the metabolites that were completely labeled by ^13^C, because at early time points this fraction of the pool is likely to be derived directly from the [U-^13^C]-glucose. For G6P, this means all six carbons are ^13^C (M+6), while for pyruvate it means all three (M+3). For UDP-glucose, while there are 15 carbons, we considered M+6 UDP-glucose to be “ubiquitously” labeled, as at early time points these carbons likely make up the glucose moiety.

AKT inhibition decreased the fraction of the metabolite pool of G6P and UDP-glucose ubiquitously labeled by ^13^C within 10 minutes of ^13^C-glucose addition (Fig 2c). This indicated that AKT signaling either increases glucose contribution to G6P and UDP-glucose or decreases gluconeogenic or glycogenolytic contribution that would otherwise dilute out the ^13^C label. To discern which pathway was responsible, we queried the fractional contributions of other isotopologues (Fig. 2d) and found that AKT inhibition decreased M+6 G6P and UDP-glucose and M+3 glycerol 3-phosphate (G3P).

Conversely, it increased the fractional contribution of unlabeled glucose to the G6P, UDP-glucose and G3P pools and partially labeled (M+3) contribution to G6P (Fig. 2d). These data indicate that AKT could be increasing the glycolytic contribution to these metabolites and suppressing contribution from the indirect pathway. Labeling of glycolytic intermediates and products downstream of G6P such as pyruvate and lactate were not consistently, significantly perturbed by MK2206 (Fig. 2d). Looking at total ion counts (Fig. 2e), differences in the ubiquitously labeled isotopologues for G6P, UDP-glucose, and G3P largely drove the differences in fractional contributions (Fig. 2d). These results are consistent with the model that AKT signaling rapidly promotes glucose phosphorylation and commitment to glycogenic precursors in hepatocytes.

Interestingly, AKT inhibition did not affect the total pool size of G6P or UDP-glucose (Fig. 2e). This indicates that the hepatocyte maintains its total pool of G6P and UDP-glucose, such that a concomitant increase in endogenous sources of carbon, or decrease in G6P consumption, compensate if glucose contribution decreases. G3P is an interesting exception, as the total pool size of G3P was reduced at multiple time points by a loss in AKT signaling (Fig. 2e), suggesting that its metabolism is controlled by AKT through a distinct mechanism.

### 3.3. Acute AKT inhibition increases the contribution of endogenous carbon sources to glucose 6-phosphate and intracellular glucose

Given that G6P and UDP-glucose pool sizes were maintained with MK2206 treatment, we explored whether endogenous sources of carbon were compensating for the decrease in glucose contribution (Fig 3a-c). M+1, M+2 and M+3 G6P and glucose can be generated from [U-^13^C]-glucose being metabolized through some combination of the pentose phosphate pathway, glycolysis, and the citric acid cycle, and then combining with endogenous carbon sources to re-generate G6P and glucose through gluconeogenesis. Notably, the [U-^13^C]-glucose tracer is not 100 percent M+6 glucose, as we detected two percent M+5 and .7 percent M+4 glucose (data not shown). The M+4 and M+5 fractions of intracellular and media glucose were not enriched compared to the glucose tracer, so alternate sources of their generation were not considered.

**Figure 3.**
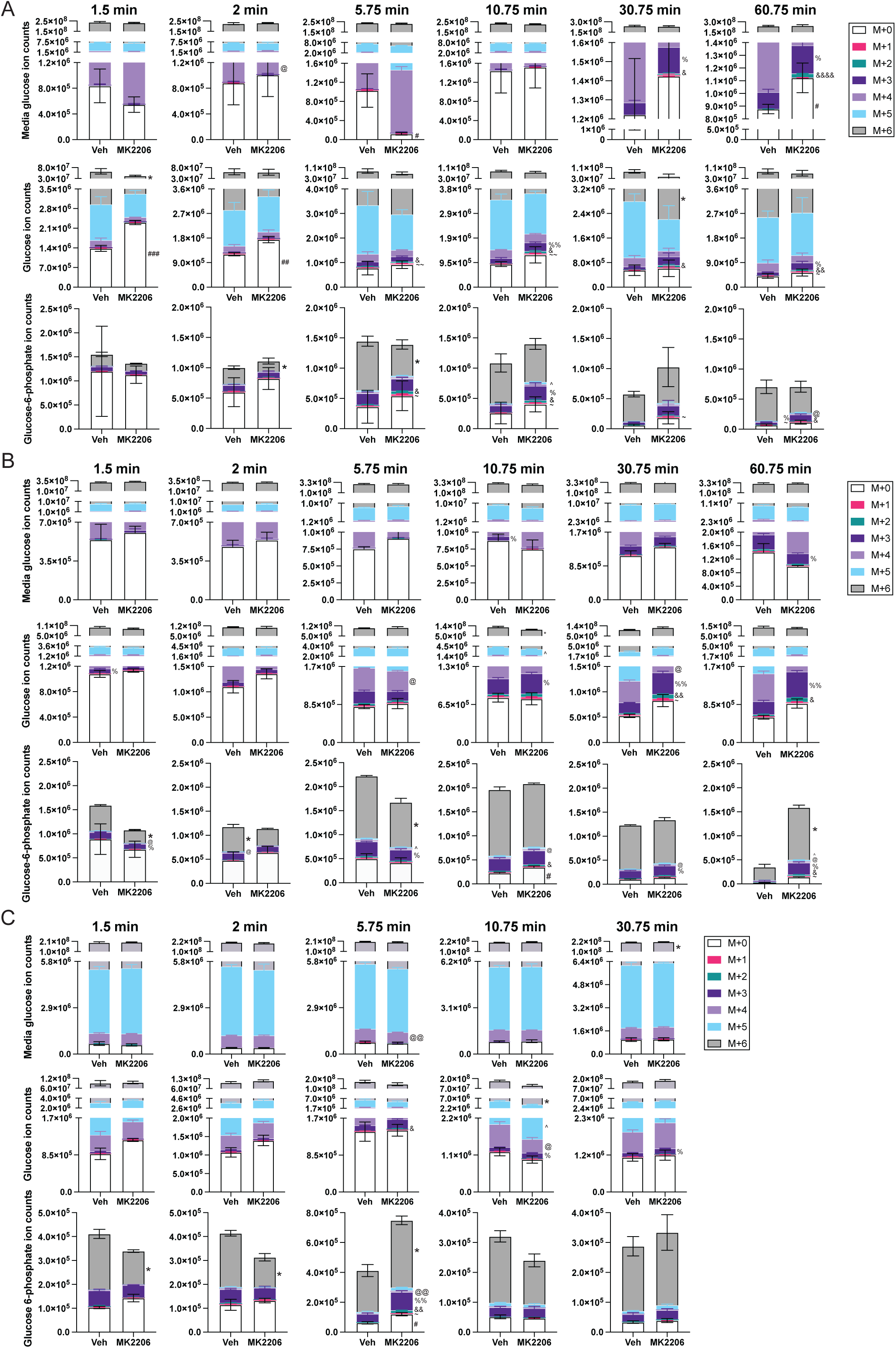
Acute AKT inhibition increases the contribution of endogenous carbon sources to glucose 6-phosphate and intracellular glucose. A) Ion counts from a tracing experiment performed as described in Figure 2c-e and represented in Fig 2e. (B) and (C) are additional experimental replicates. Time point indications refer to all graphs directly underneath. N=3 replicates per treatment. P-values calculated using Student’s t-test, values <0.05 denoted by * (M+6), ^ (M+5), @ (M+4), % (M+3), & (M+2), ∼ (M+1), or # (M+0). P-values <0.005, 0.0005, or 0.00005 for any MID are denoted by one, two, or three duplicates of the symbol. Error bars represent +/- SEM. Time points indicate the time cell extraction buffer was added to wells.

In intracellular glucose (Fig. 3a-c), we consistently saw increases in M+1, M+2, and M+3 glucose ion counts, and sometimes all three, as early as 5.75 minutes. In one experiment (Fig. 3a), this was accompanied by M+0, M+2, and M+3 glucose accumulating in the media by 30.75 to 60.75 minutes with MK2206 treatment. Decreased M+6 G6P (Fig. 3a-c) always preceded the changes to M+1 through M+3 intracellular and media glucose. The appearance of M+1 through M+3 G6P and intracellular glucose between 5.75 and 30.75 minutes with MK2206 treatment suggests that AKT inhibition causes an increase in the relative contribution of endogenous carbon sources via gluconeogenesis. Because total pool sizes of metabolites affected by MK2206 are unchanged (Fig 2e, 3a-c), it follows that endogenous sources, or consumption, change to compensate for any decrease in glucose contribution to G6P or intracellular glucose, possibly via gluconeogenesis.

### 3.4. Acute AKT inhibition does not affect glycolytic or gluconeogenic enzyme protein levels or total pool sizes of glycolytic metabolites

Insulin controls glycolytic and gluconeogenic gene expression through AKT [14], specifically *Gck*, *Pck1*, and *G6pc*. Given this, we determined whether acute AKT inhibition affected gene expression and protein levels to drive glucose contribution to G6P and glycogenic intermediates. We treated primary rat hepatocytes with unlabeled 10 mM glucose in the presence or absence of MK2206 for 30 minutes, using the same experimental paradigm as before (Fig. 2). AKT inhibition did not significantly affect expression (Fig. 4a) or total protein for GCK, PEPCK and G6PC (Fig. 4b), suggesting that AKT does not drive glucose phosphorylation through a change in GCK or G6PC protein levels.

**Figure 4.**
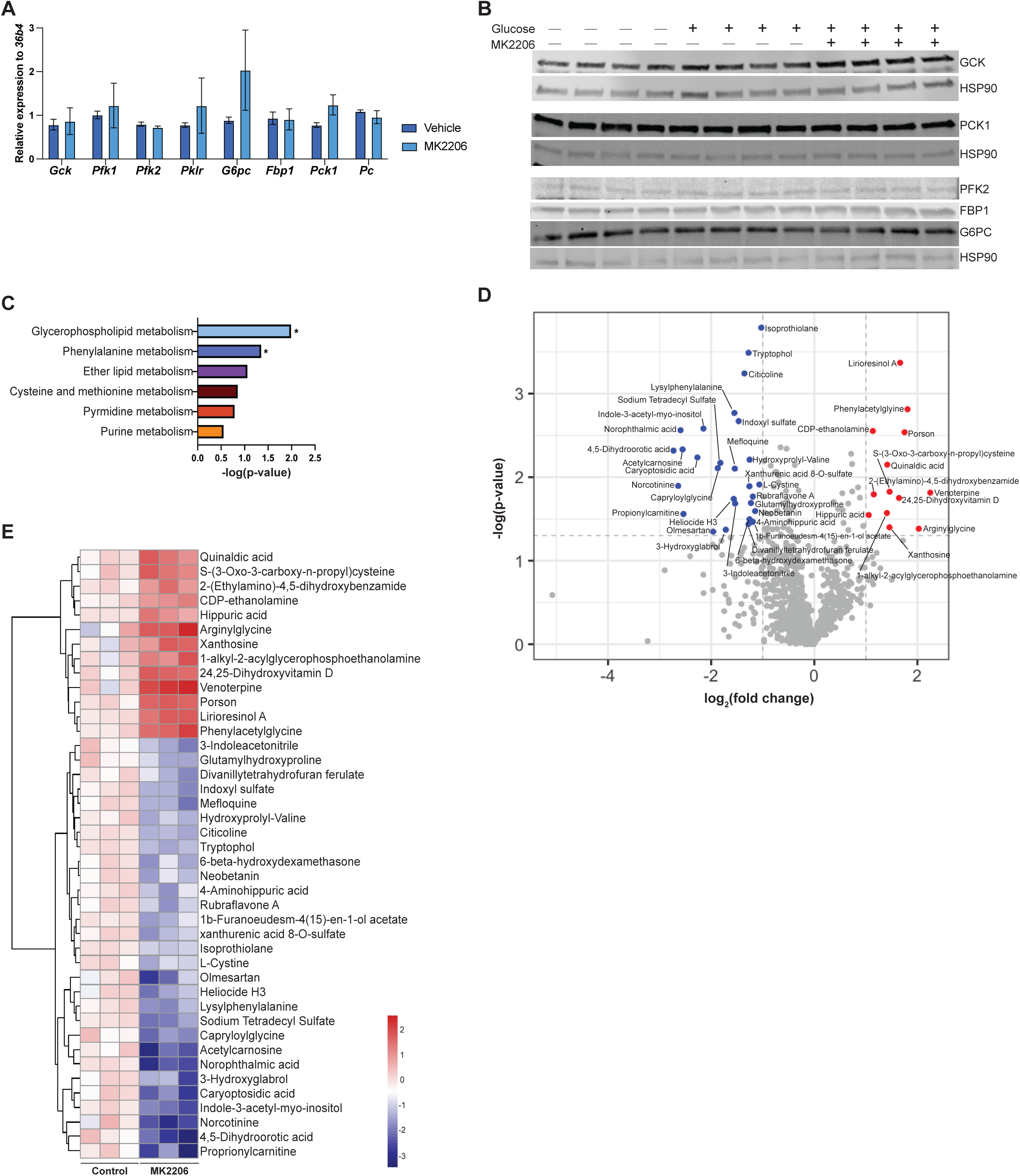
Acute AKT inhibition does not affect glycolytic or gluconeogenic enzyme protein levels or total pool sizes of glycolytic metabolites. A) Relative RNA expression for glycolytic and gluconeogenic enzymes in primary rat hepatocytes incubated overnight in serum and glucose-free DMEM, pretreated for 30 minutes with 10 μM MK2206 or DMSO in overnight media, then treated with 10 mM glucose with or without 10 μM MK2206 or DMSO for 30 minutes. Housekeeping gene is acidic ribosomal phosphoprotein P0 (*36b4*). *Gck*: glucokinase, *Pfk1:* phosphofructokinase 1, *Pfk2*: 6-phosphofructo-2-kinase/fructose-2,6-bisphosphatase, *Pklr*: pyruvate kinase, *G6pc:* glucose 6-phosphatase, *Fbp1*: fructose-1,6-bisphosphatase 1, *Pck1*: phosphoenolpyruvate carboxykinase 1, *Pc:* pyruvate carboxylase. N=4 experimental replicates averaged from 2-4 technical replicates. Error bars represent +/- SEM. B) Protein levels of glycolytic and gluconeogenic enzymes in primary rat hepatocytes treated as described in (A). HSP90 as a loading control corresponds to the blots directly above it. C) KEGG pathway analysis performed on metabolites identified through bulk metabolomics on primary rat hepatocytes treated with MK2206 for five minutes. Cells were fasted overnight in serum and glucose-free DMEM and then pretreated for 30 minutes with 10 µM MK2206 or DMSO. Glucose (10 mM) and 10 µM MK2206 or DMSO was then added to the cells for five minutes before metabolism was quenched and lysates were collected. *p-value<0.05, enrichment determined using Fisher’s exact test. D) Volcano plot and E) individual metabolites showing metabolites significantly up- or downregulated by an average fold-change +/- 2 with a p-value <0.05 in experiment described in (C). N=three samples per group. Legend in (E) is log2(fold change).

Next, given the differential effects of AKT activity on total pool sizes (Fig. 2e), we wanted to explore the global effect of acute AKT signaling on hepatocyte metabolism. To this end, we conducted unbiased metabolomics on primary hepatocytes after five minutes’ exposure to 10 mM glucose and MK2206. Cells were treated exactly as described in Figure 2. The top pathway affected was glycerophospholipid metabolism followed by phenylalanine metabolism (Fig. 4c). There were trends in enrichment of other lipid and amino acid metabolic pathways, namely ether lipid metabolism, which can be impacted downstream of glycerophospholipid metabolism (Fig. 4c). In contrast to the metabolomics conducted on AKT knockout livers (Fig. 1c), there was no trend towards enrichment of glycolysis or gluconeogenesis.

In evaluating endogenous metabolites impacted by inhibition of AKT, phospholipid precursors stood out. Specifically, CDP-ethanolamine was increased relative to controls and citicoline (CDP-choline) was decreased (Fig. 4d-e). Each of these compounds contribute to distinct arms of the Kennedy pathway to generate phosphatidylethanolamine (PE) and phosphatidylcholine (PC), respectively. This suggests that AKT may acutely downregulate PE synthesis and favor PC through a post-translational mechanism. Additionally, 1-alkyl-2-acylglycerophosphoethanolamine was also elevated with AKT inhibition (Fig. 4d-e), which is proposed to be a substrate of ethanolamine phosphotransferase (EPT1), an enzyme that generates PE from 1,2-diacylglycerol (DAG) and CDP-ethanolamine [31]. Of note, the total levels of G3P, a precursor of phosphatidylcholine synthesis, trended down in MK2206-treated cells (p-value = 0.064, data not shown), consistent with the reduction in total ion counts observed with stable isotope tracing (Fig. 2e).

Notably, glucose 6-phosphate abundance was not impacted by AKT inhibition, consistent with our stable isotope tracing experiments (Fig. 2e). This is further evidence that the reduction in G6P observed in AKT1/2 knockout livers is due to the chronic loss of GCK transcription [12] and translation [32]. Overall, short-term inhibition of AKT signaling does not affect abundance of glycolytic intermediates. The differences between these results and that of our targeted, isotope tracing approach highlights the ways in which bulk metabolomics conceals dynamic changes in metabolic pathways.

### 3.5. AKT directs glucose contribution to glucose 6-phosphate, UDP-glucose and glycogen synthesis independent of glycogen breakdown

Our stable isotope tracing data (Fig. 2) showed that AKT promotes the fractional contribution of glucose to G6P and UDP-glucose, and this preceded significant changes to endogenous sources of carbon. One way in which this could occur is by decreasing glycogenolysis to prevent the appearance of unlabeled G6P. To test this hypothesis, we evaluated [U-^13^C]-glucose metabolism in the presence of a glycogen phosphorylase inhibitor. Glycogenolysis accounts for about 45% of glucose production during the first 24 hours of fasting in humans [33] and an estimated two-thirds of HGP in conscious dogs [7]. Insulin suppresses glycogen breakdown [34] and promotes its storage [35].

AKT controls both glycogen synthesis and breakdown through transcriptional and post-translational mechanisms [17; 36; 37], and there is no evidence that AKT regulates gluconeogenesis acutely, so we tested the hypothesis that AKT promotes glucose contribution to G6P and UDP-glucose through suppression of glycogen breakdown.

We treated cells with Glycogen Phosphorylase Inhibitor (GPI) in the presence or absence of MK2206, using the same experimental approach as in Figure 2. We found that treatment with GPI alone did not significantly increase the fractional contribution of ubiquitously labeled G6P or UDP-glucose to the total pool size, nor did it rescue the effect of AKT inhibition (Fig. 5a). Investigating the contribution of other isotopologues (Fig. 5b-c), we found that AKT inhibition decreased [U-^13^C]-glucose contribution to G6P and UDP-glucose pools regardless of glycogen breakdown inhibition. Because GPI had minimal effect, we tested its efficacy in primary hepatocytes and found that it increased glycogen levels as expected when compared to glucose alone (Fig. 5d). Overall, these data suggest that AKT rapidly promotes glucose contribution to G6P and UDP-glucose through a mechanism independent of glycogen breakdown.

**Figure 5.**
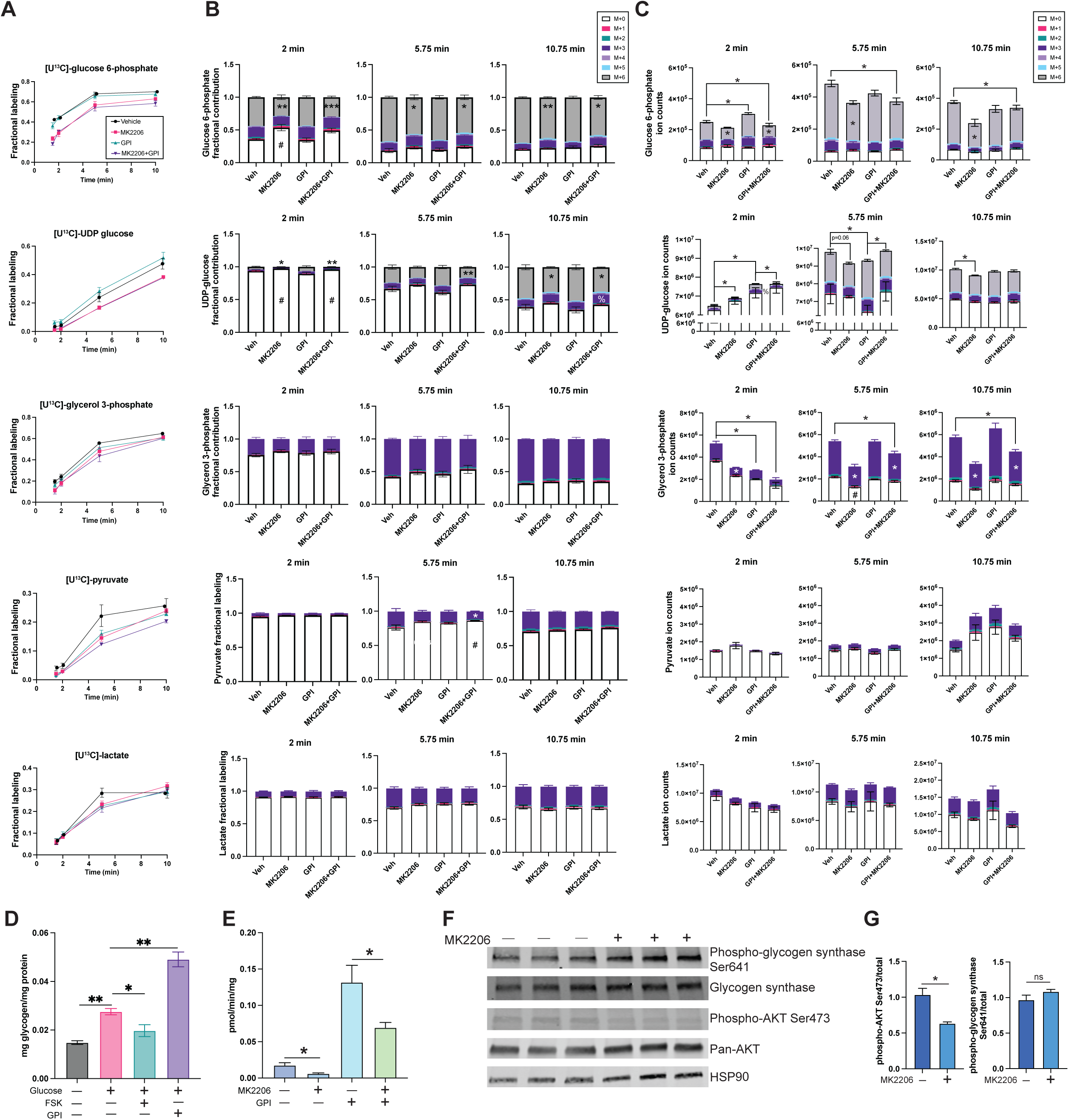
AKT directs glucose contribution to glucose 6-phosphate, UDP-glucose and glycogen synthesis independent of glycogen breakdown. A) Fractional labeling and B) fractional contribution of metabolite pools in primary rat hepatocytes treated with [U-^13^C]-glucose with or without 10 µM MK2206 and 5 µM GPI. Cells were treated as described in Section 2.2. N=six replicates from two pooled experiments. C) Ion counts from a representative tracing experiment included in the analysis done in (A) and (B). P-values calculated using a one-way ANOVA followed by a post-hoc t-test, values <0.05 denoted by *(M+6), %(M+3), or # (M+0). Values <0.005 and <0.0005 for M+6 denoted by ** and ***. P-values on top of, or adjacent to, specific isotopologues compare MK2206 and Veh, or GPI+MK2206 and GPI. Time points indicate the time cell extraction buffer was added to wells. D) Total glycogen content in primary rat hepatocytes following treatment with 20 mM glucose with or without 25 μM FSK and 5 μM GPI for four hours. P-values calculated using one-way ANOVA and post-hoc t-test, *p-value <0.05, **p-value<0.005. E) Glycogen synthesis in primary rat hepatocytes that were fasted overnight in serum and glucose-free media, pretreated for 30 minutes with 10 μM MK2206 or DMSO in overnight media, and then treated with 1 µCi [U^14^C]-glucose and 10 mM cold glucose with inhibitors or DMSO for 30 minutes. N=four to six replicates pooled from five experiments. *P-value<0.05, calculated using a one-way ANOVA followed by a post-hoc t-test. F) Western blot of glycogen synthase phosphorylation at Ser641 in primary rat hepatocytes incubated overnight in serum and glucose-free DMEM, pretreated for 30 minutes in overnight media with 10 μM MK2206 or DMSO, and then treated with 10mM glucose with inhibitor or DMSO for 30 minutes. G) Quantification for blots shown in (F), phospho-antibody signal normalized to its respective total protein. *P-value<0.05 calculated using Student’s t-test. All error bars represent +/- SEM.

Our results from stable isotope tracing (Fig. 2,5) show a decrease in the synthesis of G6P and UDP-glucose from glucose with AKT inhibition, indicating that there may be a decrease in glycogen synthesis through the direct pathway. To determine the effect of acute AKT inhibition on macromolecule synthesis, we measured glycogen synthesis using the same experimental paradigm as in Figure 5a-c but with [U-^14^C]-glucose. We found that AKT inhibition decreased glucose contribution to glycogen following a 30-minute incubation with the radiolabel (Fig. 5e), too rapid to be accounted for by a transcriptional mechanism. Of note, GPI treatment alone increased glycogen synthesis dramatically, highlighting that significant glycogen cycling occurs in primary hepatocyte culture. Nevertheless, MK2206 treatment suppressed glycogen synthesis in the presence of GPI (Fig. 5e), demonstrating that AKT rapidly promotes glucose contribution to glycogen independent of inhibiting its breakdown.

There are several known mechanisms that could potentially explain how AKT promotes glycogen synthesis post-translationally. G6P allosterically activates glycogen synthase, increasing glycogen storage [22; 23; 38]. However, our unbiased metabolomics and stable isotope tracing data show that while glucose contribution to G6P is affected by AKT inhibition, the total pool size is unchanged, making this an unlikely explanation.

Insulin signaling through AKT can also control glycogen synthesis through reversible phosphorylation of glycogen synthase. AKT inhibits GSK-3β [17], which phosphorylates glycogen synthase to inhibit it [39]. Second, AKT can activate the PPP1R3G subunit of the Protein Phosphatase 1 (PP1) complex [36], stimulating its activating dephosphorylation of glycogen synthase and inhibitory dephosphorylation of glycogen phosphorylase [40]. To determine if AKT increases glucose contribution to glycogen on this short timescale through control of glycogen synthase phosphorylation, we blotted our cell lysates for glycogen synthase Ser641 phosphorylation, which is associated with its inhibition [41–44]. We found that AKT inhibition failed to increase Ser641 phosphorylation (Fig. 5f-g), indicating that the decrease in glycogen synthesis could be occurring independent of glycogen synthase. This is consistent with previous reports that mutation of AKT-regulated phosphosites on GSK-3β does not impact glycogen synthesis *in vivo* [18], suggesting additional pathways are responsible. The observed decrease in glycogen synthesis from glucose with acute AKT inhibition may be a potential consequence of the upstream defect in glucose contribution to G6P and UDP-glucose.

## 4. Discussion

Insulin’s control of HGP and stimulation of liver glucose uptake is well appreciated[4]. Previous studies suggest involvement of hepatic AKT in insulin-mediated control of HGP, whether through suppressing glycogenolysis or gluconeogenesis [14; 17; 36]. Data in this manuscript suggest that AKT signaling may also stimulate glycogen synthesis through the direct pathway, that is, from extracellular glucose and not gluconeogenic intermediates, by increasing glucose contribution to G6P. Previous studies have focused on the importance of HGP suppression for insulin-stimulated liver glucose uptake, specifically inhibition of glycogenolysis. Data presented here suggest an additional mechanism independent of glycogen breakdown that may precede eventual changes to endogenous glucose production. Broadly, our study reveals an alternative to a substrate push mechanism, whereby AKT activation increases glucose phosphorylation while decreasing contribution from endogenous carbon sources to increase glucose contribution to UDP-glucose and glycogen without increasing total levels of G6P or UDP-glucose.

The most robust effect of AKT signaling was on glucose contribution to G6P and UDP-glucose, as indicated by changes in the ubiquitously labeled fraction of each metabolite (Fig. 2d and 5b). At certain time points, there were increases in contribution of non-ubiquitously labeled G6P, indicating increased flux through gluconeogenesis (Fig 2d, 3a-c, 5b). A substrate push mechanism may be appropriate to explain the effect of MK2206 on endogenous carbon contribution to G6P, whereby transient decreases in the G6P pool caused by decreased glucose phosphorylation could drive an increase in its generation from gluconeogenic precursors. These potential transient changes, however, do not persist long enough to lead to significant differences in pool size, making G6P allosteric activation of glycogen synthase an unlikely explanation for MK2206’s eventual effect on glycogen synthesis (Fig. 5e).

An advantage of stable isotope tracing compared to earlier work done with radioisotopes is that it allows a more complete network picture of insulin’s control of glucose metabolism. Earlier studies conducting clamps in the conscious dog model concluded that insulin did not significantly affect gluconeogenic flux [19; 20], but authors acknowledged that experimental pitfalls could preclude accurate measurement. First, while insulin might suppress gluconeogenesis, this could be masked by its dominant effect on glycogen breakdown [6], whereby decreased G6P due to decreased glycogenolysis leads to an increase in gluconeogenesis, which would appear as unchanged flux. Second, experiments estimating gluconeogenic flux using ^14^C-phosphoenolpyruvate or ^14^C-lactate and measuring incorporation into ^14^C-glucose in the plasma may underestimate changes in gluconeogenic flux if insulin diverts ^14^C-G6P derived from HGP to glycogen synthesis instead of glucose export. Third, several studies raised skepticism regarding the accuracy of using ^2^H_2_O incorporation into glucose to measure gluconeogenesis vs glycogenolysis, as glyceraldehyde 3-phosphate (GAP) exchange with fructose 1,6-bisphosphate or retention of a deuterium during conversion of GAP to dihydroxyacetone phosphate (DHAP) can lead to underestimation of gluconeogenesis [45; 46]. Using our approach, we were able to determine dilution of our tracer with or without glycogen breakdown to approximate contribution of gluconeogenesis. We also demonstrate that insulin signaling, at least through AKT, does indeed divert G6P to glycogen rather than downstream to glycolysis.

Considerable effort has been made to quantify insulin’s control of gluconeogenesis [6; 15; 19-21]. It has been proposed that gluconeogenic flux does not change significantly even with hyperglycemia [47; 48], and that perhaps only fatty acid flux controls rates of gluconeogenesis [6; 49]. Consistent with this notion, Basu *et al.* showed that type 2 diabetes mellitus patients have elevated glycogenolysis but unchanged gluconeogenesis in response to a hyperinsulinemic clamp compared to mild diabetics or non-diabetic controls [11]. Together, the literature suggest that insulin suppresses gluconeogenesis through inhibition of adipose tissue lipolysis and not through direct action at the liver. Our observations (Fig. 3a-c) suggest that insulin signaling via AKT may indirectly decrease gluconeogenesis secondary to promoting glucose phosphorylation. Interestingly, the increased M+1, M+2, and M+3 intracellular glucose in our experiments (Fig. 3a-c) was not at equilibrium with the glucose in the media, as those isotopologues did not become enriched in media glucose until at least 30 minutes. This indicates that intracellular glucose may become re-phosphorylated more readily than exported in primary hepatocytes.

Our data also sheds light on whether a “pull” mechanism governs insulin’s control of glycogen deposition. AKT inhibition decreased glucose contribution to G6P, UDP-glucose, and glycogen synthesis (Fig. 2d-e, 5a-c, 5e), evoking the question of whether decreased glycogen synthesis created less of a demand for glucose to contribute to the generation of glycogenic metabolites. However, if changing glycogen levels influence glucose contribution to G6P and UDP-glucose, M+6 labeling of G6P would increase when glycogen breakdown is inhibited. Instead, GPI had a strong effect on glycogen synthesis (Fig. 5e) with little effect on glucose labeling of G6P beyond 2 minutes (Fig. 5b-c), relative to the control. This suggests that the effect of AKT inhibition on glucose contribution to G6P is independent of glycogen breakdown and that glucose phosphorylation is rapid relative to the rate of glycogen synthesis.

Together, these data suggest the possibility of a novel mechanism where AKT acutely promotes extracellular glucose contribution to glycogen synthesis by rapidly increasing glucose phosphorylation. Previous work proposed that AKT increases glycogen synthesis post-translationally by indirectly controlling phosphorylation of glycogen synthase [17], although we did not detect a significant increase in glycogen synthase phosphorylation in isolated rat hepatocytes (Fig. 5f-g). Additionally, while MK2206 decreased glycogen synthesis in the presence of GPI, glycogen synthesis in the combination group was elevated compared to MK2206 treatment alone (Fig. 5e). The increase in glycogen synthesis when glycogen breakdown is inhibited demonstrates that there is sufficient activity of glycogen synthase under these conditions. If MK2206’s effect on glycogen synthesis from glucose depended on glycogen synthase, GPI treatment with MK2206 would fail to increase glycogen synthesis relative to the control or MK2206 alone.

With regard to the mechanism by which AKT acutely promotes glucose contribution to G6P and then to glycogen, the effect of MK2206 on upper glycolysis implicates a change in GCK activity (Fig 2c-e, Fig 5a-c). This is consistent with previous work demonstrating that insulin activates GCK [24; 26; 50; 51]. Our results are also consistent with a report showing that postprandial GCK activator treatment in rats increases glycogen deposition [52]. G6Pase overexpression, which reduces G6P levels, reduces glycogen synthesis and glycolysis [53], while GCK overexpression increases G6P, glycogen deposition, and total levels of glycolytic intermediates [54]. How AKT might be activating GCK is unclear, but it is well-known that GCK activation is regulated through its subcellular localization. During fasting, when intracellular glucose levels are low, GCK is sequestered in the nucleus by Glucokinase Regulatory Protein (GKRP). Rising blood glucose levels postprandially favor glucose transport into the liver. Glucose diffuses into the nucleus, binds to GKRP, and consequently releases GCK to be transported back into the cytoplasm where it can phosphorylate glucose [55]. Interestingly, we did not see a change in GCK localization in primary rat hepatocytes treated with 10 mM glucose with or without 10 µM MK2206 (data not shown). This suggests that if the mechanism is GCK-dependent, it does not occur through its translocation.

Beyond the short-term effects of AKT signaling on upper glycolytic metabolites, we also observed changes to glycerophospholipid metabolism. AKT inhibition increased CDP-ethanolamine and decreased CDP-choline (Fig. 4d-e), which each contribute to glycerophospholipid synthesis through distinct arms. Thus, AKT may be increasing the phosphatidylethanolamine arm of phospholipid synthesis through post-translational regulation. CCTα is the enzyme catalyzing CDP-choline synthesis and its activity is regulated by mTORC1 in the liver [56]. AKT activates mTORC1 to drive lipogenesis [12], thus it is possible that loss of mTORC1 activity due to AKT inhibition decreased CDP-choline synthesis. It is unclear why a loss in AKT activity leads to an increase in CDP-ethanolamine levels, although it may be due to a compensatory mechanism in which PE synthesis is upregulated to increase PC generation via phosphatidylethanolamine N-methyltransferase (PEMT) activity. We also observed in our stable isotope tracing experiments (Fig. 2e and 5c) that acute AKT inhibition decreased G3P levels. This was recapitulated in our bulk metabolomics experiments, where G3P trended to be reduced (p = 0.06, data not shown). G3P is acylated by the GPAT enzymes in the DAG synthesis pathway, which insulin has been shown to activate in adipocytes [57]. However, the isoform of GPAT1 expressed in hepatocytes does not contain an AKT consensus phosphorylation motif, so it is unlikely that G3P contribution to DAG synthesis was increased by acute AKT inhibition. G3P is synthesized from DHAP by GPD1, but GPD1 also does not contain an AKT consensus phosphorylation motif. Thus, future studies are needed to investigate how AKT acutely controls G3P levels in hepatocytes.

With acute inhibition of AKT, we did not observe significant differences in the levels of lactate [27–29] or pyruvate [27] as previous studies had, but we did detect the difference in G3P reported by Terrettaz *et al* [27]. Discrepancies may reflect differences between insulin and AKT-specific actions, given these previous reports treated with insulin. Additionally, one study described above [28] cultured primary hepatocytes for 48 hours, at which point primary hepatocytes begin to de-differentiate. There may also be differences in the degree of insulin signaling occurring in the controls that can affect results. In our hands, even serum-starved primary rat hepatocytes exhibit residual AKT phosphorylation (Fig. 2b, lane 1), demonstrating that this pathway is difficult to suppress in culture and indeed is likely required to prevent apoptosis [58]. However, without being able to compare the level of activation of signaling intermediates to other studies’ controls, it is difficult to discern the root of discrepancies. This is a limitation of our study.

In summary, we have provided evidence that AKT acutely stimulates glucose phosphorylation and contribution to glycogen, independent of glycogen breakdown. This may be occurring independent of glycogen synthase regulation. Future studies are needed to determine the precise mechanism by which AKT is achieving this, with a glucokinase-dependent mechanism likely. Considering that decreased liver GCK expression in humans correlates strongly with worsened blood glucose control [16], and diabetes patients exhibit decreased GCK activity [59], understanding this mechanism likely holds implications for treating hyperglycemia.

## Author contributions

MLS: Conceptualization, Investigation, Methodology, Formal Analysis, Funding Acquisition, Writing-Original Draft. WDL and Rabinowitz Lab: Methodology, Formal Analysis. TC: Investigation, Formal Analysis. JB and JDR: Conceptualization, Writing-Review & Editing. PMT: Conceptualization, Investigation, Formal Analysis, Writing-Review & Editing, Supervision, Project Administration, Funding Acquisition.

## Acknowledgements

We thank the Penn Vector Core, Penn Diabetes Research Center (P30-DK19525), and the Penn Center for Molecular Studies in Digestive and Liver Diseases (P30-DK050306). We also thank all the members of the Titchenell and Baur labs for their feedback and fruitful discussions on this project.

This work was supported by U.S. National Institutes of Health (NIH) grant R01-DK125497 (PMT), internal funds from the University of Pennsylvania (P.M.T.), NIH NRSA F31DK139617 (MLS), NIH NRSA F32-DK127843 (WDL), and Cox Research Institute.

